# ARF suppresses 5’-terminal oligopyrimidine mRNA translation

**DOI:** 10.1101/2020.02.07.939165

**Authors:** Kyle A. Cottrell, Ryan C. Chiou, Jason D. Weber

**Affiliations:** Department of Medicine, Division of Molecular Oncology, Washington University School of Medicine, Saint Louis, Missouri, USA; Department of Cell Biology and Physiology, Siteman Cancer Center, Washington University School of Medicine, Saint Louis, Missouri, USA

## Abstract

Tumor cells require nominal increases in protein synthesis in order to maintain high proliferation rates. As such, tumor cells must acquire enhanced ribosome production. How many of the mutations in tumor cells ultimately achieve this aberrant production is largely unknown. The gene encoding ARF is the most commonly deleted gene in human cancer. ARF plays a significant role in regulating ribosomal RNA synthesis and processing, ribosome export into the cytoplasm, and global protein synthesis. Utilizing ribosome profiling, we show that ARF is a major suppressor of 5’-terminal oligopyrimidine mRNA translation. Genes with increased translational efficiency following loss of ARF include many ribosomal proteins and translation factors. Knockout of *p53* caused a similar increase in 5’-TOP mRNA translation. The 5’-TOP regulators mTORC1, eIF4G1 and LARP1 are dysregulated in ARF and p53 null cells.

## Introduction

Accelerated cellular division and macromolecular growth of tumors is dependent on robust protein synthesis. To meet this need, ribosome biogenesis and translation rates are frequently elevated in cancer cells ^1^. Increased translation in cancer cells is often driven by activation of the mTORC1 pathway ^1^.

Loss of the gene encoding the ARF tumor suppressor is the most common copy number variation in cancer ^2^. ARF is expressed in response to oncogenic stimuli – including the overexpression of Myc, oncogenic Ras and chronic activation of the mTORC1-pathway ^3,4^. The canonical function of ARF is to stabilize p53 by sequestering MDM2 in the nucleolus ^4–6^. In addition to its canonical role, ARF suppresses global protein synthesis and ribosome biogenesis by regulating rRNA transcription and processing ^7–16^. ARF also regulates the translation of specific mRNAs, including VEGFA and DROSHA ^10,17^.

While ARF is known to regulate ribosome biogenesis, the p53-MDM2 axis senses dysregulation of ribosome biogenesis ^18^. Free ribosomal proteins, such as RPL11, bind to MDM2 and lead to stabilization of p53 ^18^. In addition, p53 is known to repress the activity of mTORC1 during times of genotoxic stress ^19^.

The synthesis of ribosomal proteins is tightly regulated by the mTORC1 pathway ^20^. Many ribosomal proteins and some translation factors contain a 5’-terminal oligopyrimidine (5’-TOP) motif at the beginning of their mRNAs ^21,22^. The translation of 5’-TOP mRNAs is regulated by LARP1 which inhibits translation through disrupting eIF4G1 binding to the 5’ end of the mRNA ^23–25^. Activation of the mTORC1 pathway leads to inhibition of LARP1 via phosphorylation by AKT and S6-kinase ^23^. When phosphorylated, LARP1 binds the 3’UTR of 5’-TOP mRNAs and enhances translation ^23^.

Here, we show that ARF selectively regulates the synthesis of ribosomal proteins and translation factors containing 5’-TOP motifs within their mRNAs. Knockdown or knockout of ARF caused increase translation of many 5’-TOP mRNAs. This effect of ARF-loss was dependent on p53 expression. Knockout of *p53* caused a similar increase in the expression of some 5’-TOP mRNA encoded proteins. Finally, we observed dysregulation of many regulators of 5’-TOP mRNA translation.

## Results

### Increased protein synthesis following loss of ARF

ARF has been previously identified as a regulator of global protein synthesis ^10,16^. To confirm these previous findings, we assessed global protein synthesis using wildtype (WT) and *Arf*^-/-^ mouse embryonic fibroblasts (MEFs). Polysome profiling of WT and *Arf*^-/-^ MEFs revealed a shift of rRNA mass towards polysomes in *Arf*^-/-^, indicating more translation in those cells, Figure 1a. This result is consistent with previous studies ^10,16^. We further confirmed this observation by measuring the rate of puromycin incorporation into nascent peptides in WT and *Arf*^-/-^ MEFs. Puromycin is a translation elongation inhibitor that is incorporated into nascent peptide during protein synthesis ^26^. Using a puromycin antibody it is possible to detect puromyclylated peptides. We observed an increase in puromycylated proteins in *Arf*^-/-^ MEFs compared to WT MEFs, Figure 1b, further confirming elevated protein synthesis in those cells.

**Figure 1:**
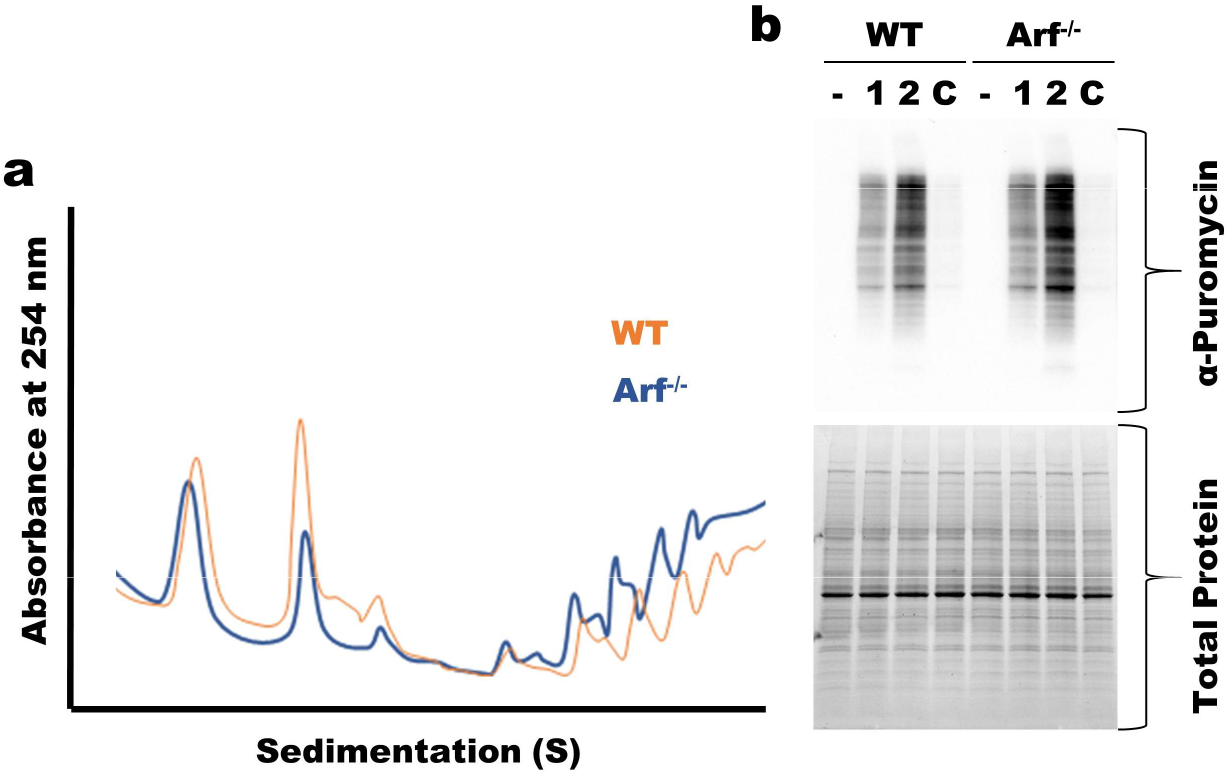
Upregulation of translation following loss of ARF. **a** Polysome profiling showing increased polysome peaks in *Arf*^-/-^ MEFs **b** Puromycin translation assay showing increased protein production in *Arf*^-/-^ MEFs. – untreated, 1 and 2 treatment with puromycin for 1 or 2 minutes, C indicates cells that were treated with cycloheximide prior to puromycin treatment for 2 minutes.

### Increased translation efficiency of 5’-TOP mRNAs upon ARF loss

Previously ARF has been identified to not only regulate translation globally but to specifically regulate the translation of a subset of mRNAs ^10,17^. To map global translational regulation by ARF we employed ribosome profiling to measure changes in translation efficiency following ARF-loss. We performed ribosome profiling and RNA-seq for WT MEFs transduced with either shSCR or shARF in addition to *Arf*^-/-^ MEFs transduced with shSCR. We identified differentially translated mRNAs using the metric translation efficiency (TE), which is the ratio of ribosome profiling reads to total RNAseq reads for each gene ^27^. Most mRNAs show no significant difference in TE upon loss of ARF, Figure 2b and c. However, a subset of mRNAs showed a slight but significant increase in translation following ARF-loss by either knockdown with shARF or knockout of *Arf*. There is a significant correlation between fold change of TE following *ARf* knockdown and knockout, Figure 2d. Gene ontology analysis of the genes with increased TE following knockout of *Arf* identified multiple GO terms associated with translation, Figure 2e. A closer look at the genes with the biggest increase in TE revealed many ribosomal proteins and translation factors. It has been well established that most ribosomal proteins and many translation factors contain a 5’-TOP motif ^21,22^. Analysis of our ribosome profiling results revealed that many of the mRNAs with increased TE are known to contain a 5’-TOP motif, Figure 2f ^21^. Furthermore, mRNAs known to contain a 5’-TOP motif show a general increase in TE following knockout of *Arf*, Figure 2g.

**Figure 2:**
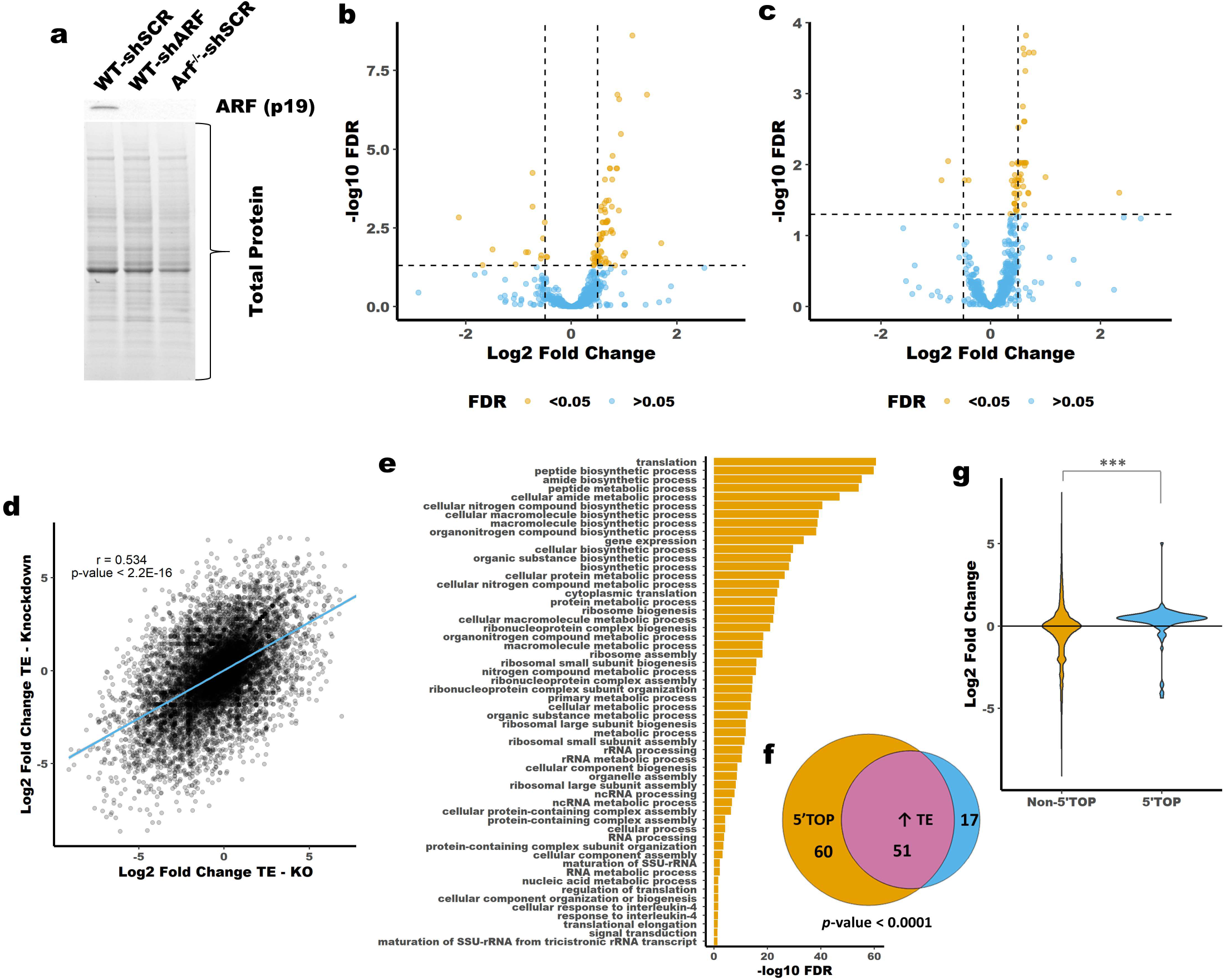
Ribosome profiling reveals upregulation of 5’-TOP mRNA translation following loss of ARF. **a** Immunoblot showing knockdown and knockout of *Arf* in MEFs. **b** Volcano plot showing Log2 Fold Change of TE between WT-shSCR and *ARF*^/-^-shSCR MEFs or WT-shSCR and WT-shARF MEFs **c**. Vertical lines are Log2 Fold Change of ±0.5. Horizontal line is FDR corrected p-value of 0.05. **d** Correlation between Fold Change of TE following ARF knockout (x-axis) or ARF knockdown (y-axis). **e** Geno ontology terms associated with genes that have increased TE following ARF knockout. **f** Venn-diagram showing overlap between mRNAs with increased TE and the presence of a 5’-TOP motif. **g** Violin plot showing the TE of mRNAs known to contain a 5’-TOP motif versus those that are not known to contain the motif ^21^. *** *p*-value <0.00001; chi-squared test (**f**) or t-test (**g**).

To validate our ribosome profiling findings, we assessed the mRNA and protein abundance of several 5’-TOP genes in MEFs following ARF-loss. By immunoblot we observed increased protein abundance for the 5’-TOP genes PABP, TPT1 and RPL22, Figure 3a-b. The expression of each gene was unchanged at the mRNA level, Figure 3c. Combined with our ribosome profiling results these data support increased translation of 5’-TOP mRNAs in *Arf*-null MEFs. To further validate these findings, we generated a 5’-TOP reporter. The promoter and 5’UTR from RPL23A, a 5’-TOP containing gene that showed increased TE upon *ARF*-loss in our ribosome profiling data, was cloned upstream of firefly luciferase, Figure 3d. This reporter was transfected into WT-shSCR, WT-shARF and *Arf*^-/-^shSCR MEFs along with a plasmid for expression of *Renilla* luciferase as transfection control. We observed increased luciferase activity for our 5’-TOP reporter in *ARF*-knockdown and knockout MEFs, Figure 3e. This contrasts with a luciferase reporter driven by the SV40 promoter that lacks the 5’-TOP motif. Analysis of the mRNA abundance of the 5’-TOP reporter revealed a slight decrease in mRNA expression in *ARF*-knockdown and knockout MEFs, Figure 3f. Together these data confirm increased translation efficiency of 5’-TOP containing mRNAs following loss of ARF.

**Figure 3:**
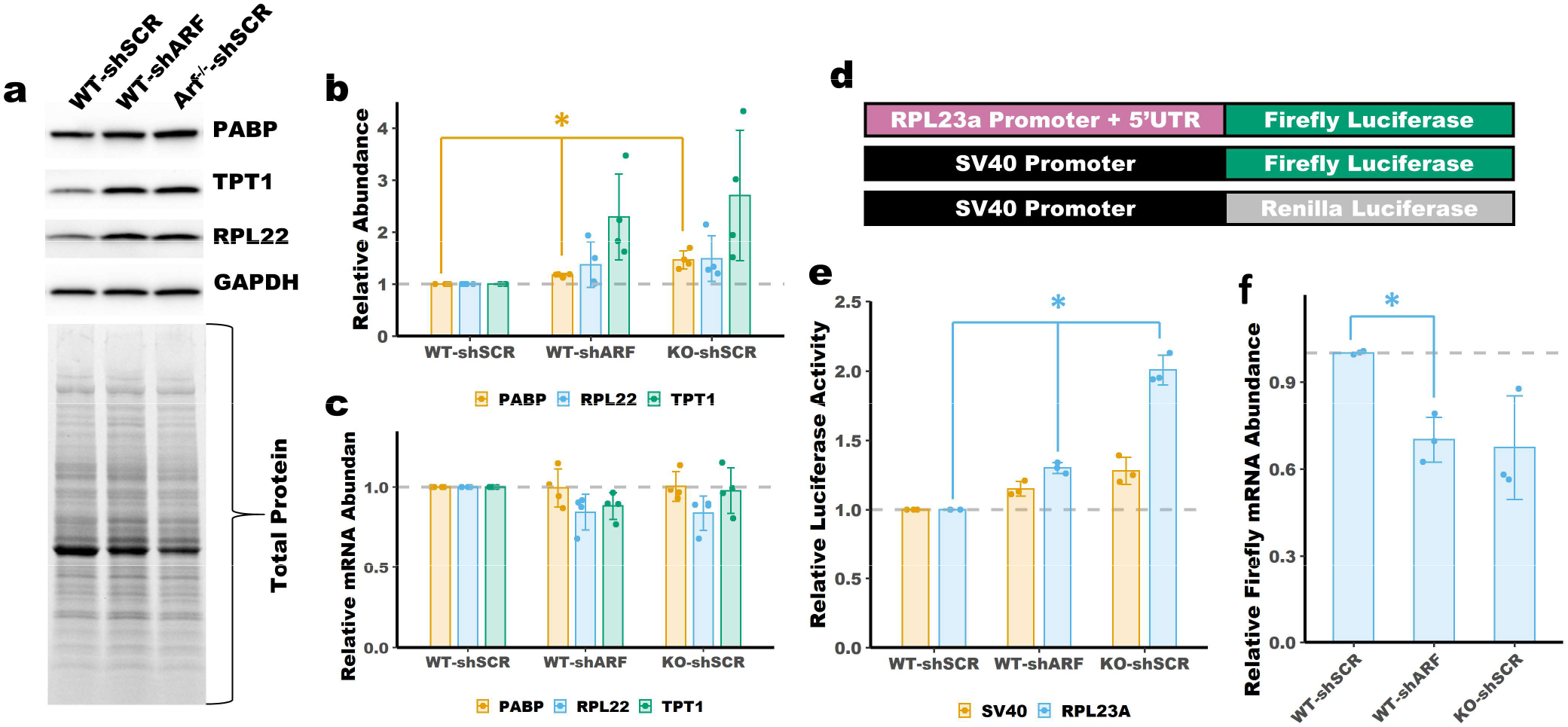
Upregulation of 5’-TOP mRNA translation following loss of ARF. **a** Immunoblot analysis showing increased protein abundance of PABP, TPT1 and RPL22. **b** Quantitation of blots in panel **a**. Total protein was used for normalization. Mean±SD, n=4. **c** qPCR shows no change in mRNA expression of 5’-TOP mRNAs. Mean±SD, n=4. **d** Schematic of the luciferase reporters used in panels **e** and **f. e** Luciferase activity of a 5’-TOP reporter is increased in ARF-null MEFs. Luciferase activity was normalized to *Renilla* luciferase transfection control and set relative to WT-shSCR. Mean±SD, n=3. **f** qPCR shows no increase e in mRNA expression of the RPL23a reporter. Normalized to *Renilla* luciferase transfection control. Mean±SD, n=3. * *p*-value < 0.05; t-test with Bonferoni correction.

### Increased translation efficiency of 5’-TOP mRNAs upon Arf loss is dependent on p53

ARF is a well-known activator of p53 ^4,5^. To test whether p53 is required for *ARF*-dependent regulation of 5’-TOP mRNA translation we performed some of the experiments described above in *p53*^-/-^ MEFs. Knockdown of *ARF* in *p53*^-/-^ MEFs had no effect on 5’-TOP mRNA protein or mRNA abundance, Figure 4a-b. Furthermore, knockdown of ARF had no effect on the translation of the RPL23A reporter, Figure 4c. There was no difference in polysome abundance between shSCR and shARF transduced *p53*^-/-^ MEFs indicating no changes in total protein synthesis, Figure 4d. However, there were some changes in the abundance 40S and 60S ribosomal subunits based on variations in peak heights. This observation mirrored that seen following knockout of *Arf* in WT MEFs. These findings indicate that p53 is required for *ARF*-dependent translational regulation of 5’-TOP mRNAs.

**Figure 4:**
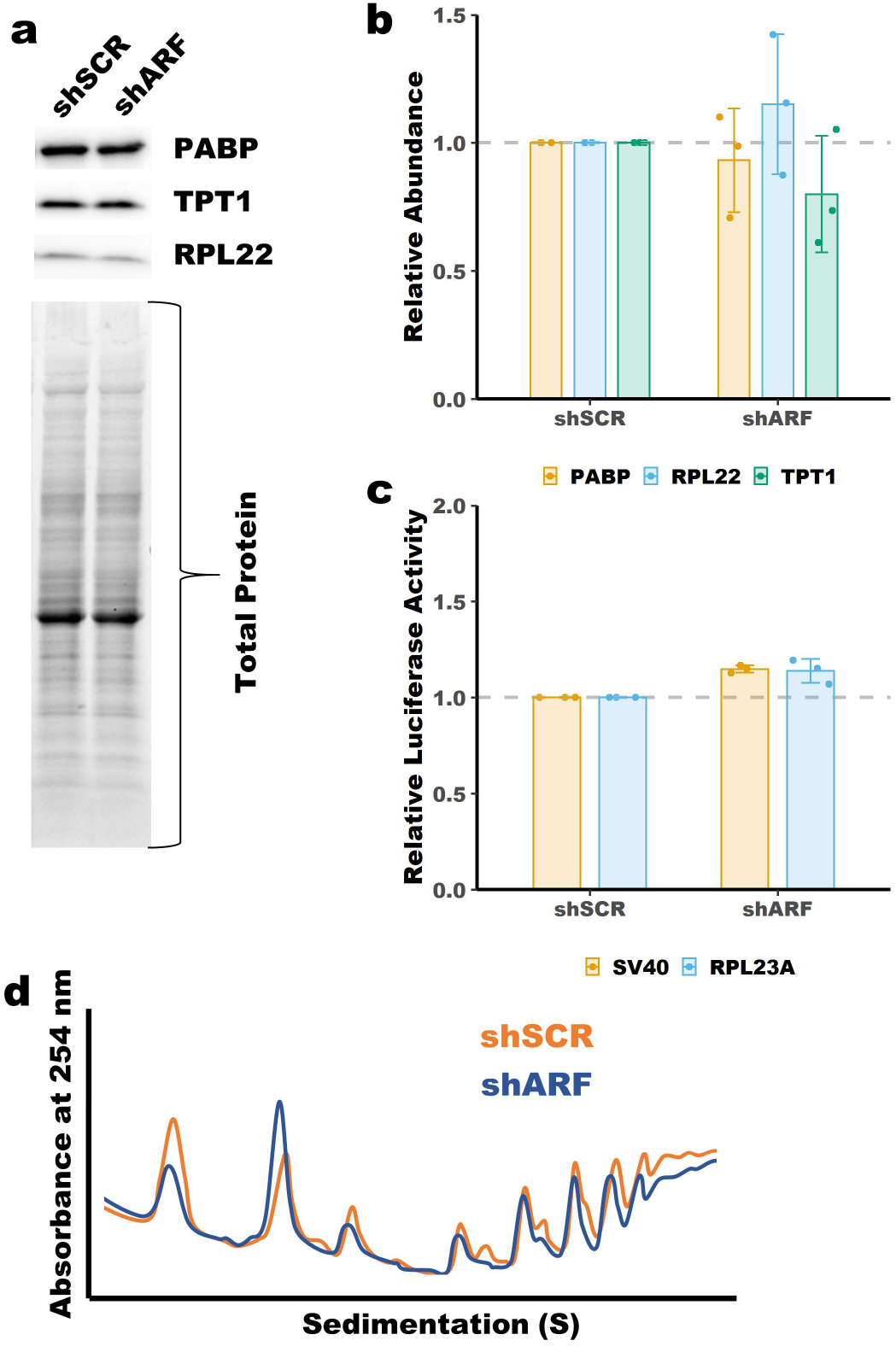
5’-TOP expression is unaffected by ARF knockdown in p53^-/-^ MEFs. **a** Immunoblot analysis showing no change in protein abundance of PABP, TPT1 and RPL22 following ARF knockdown in *p53*^-/-^ MEFs. **b** Quantitation of blots in panel **a.** Total protein was used for normalization. Mean±SD, n=3. **c** No change in luciferase activity of a 5’-TOP reporter following ARF knockdown in *p53*^-/-^ MEFs. Luciferase activity was normalized to *Renilla* luciferase transfection control and set relative to shSCR. Mean±SD, n=3. **d** Polysome profiling showing no change in polysome peaks in *p53*^-/-^ MEFs following knockdown of ARF.

### Loss of ARF modestly affects the expression or activity of 5’-TOP regulators

The mTOR pathway is known to regulate 5’-TOP mRNAs through the phosphorylation and inactivation of LARP1 ^23–25^. LARP1 is phosphorylated by the mTOR and the mTORC1 substrate S6-kinase ^23^. Immunoblot analysis of mTORC1 pathway activation in WT and ARF-null MEFs revealed a slight, though inconsistent increase in mTORC1 activity, Figure 4a-b. S6-kinase showed the largest increase, though statistically insignificant, in phosphorylation following ARF-loss, indicating LARP1 maybe phosphorylated and inactivated in those cells. However, inhibition of S6-kinase had no effect on the expression of 5’-TOP encoded proteins, Supplementary Figure 7. Interestingly, we observed a slight increase in mTORC1 pathway activation in *p53*^-/-^ MEFs following ARF-knockdown, Supplemental Figure 6. Treatment with the mTOR inhibitor Rapamycin was employed to assess the importance of mTORC1 pathway activity in *Arf*^-/-^ MEFs. Consistent with previous reports, cells treated with rapamycin show reduced mTOR autophosphorylation and reduced expression of the 5’-TOP encoded proteins RPL22, PABP and TPT1, Figure 4c-d, ^28,29^. These results show that mTORC1 activity is required for maximal expression of 5’-TOP mRNAs in ARF-null MEFs. The slight increase in mTORC1 seen following ARF knockdown or knockout could be driving increased translation of 5’-TOP mRNAs.

The 5’-TOP mRNA regulator LARP1 is known to function by competing with the translation initiation factor eIF4G ^23,25^. Immunoblot analysis revealed a slight increase in eIF4G expression in ARF-null MEFs, Figure 4e-f. The same increase was observed in *p53*^-/-^ MEFs following ARF-knockdown, Supplementary Figure 6. We attempted to overexpress eIF4G1 in WT MEFs via lentiviral transduction, but due to the size of the coding sequence the transduction efficiency was extremely low and thus we were unable to assess the importance of increased eIF4G1 expression in ARF-null MEFs (data not shown).

LARP1 is a known regulator of 5’-TOP mRNA translation ^23–25^. Immunoblot analysis of WT and ARF-null MEFs showed increase protein abundance for LARP1, Figure 5f. There was no increase in LARP1 mRNA expression in ARF-null MEFs relative to WT MEFs, Figure 5g. Knockdown of ARF in *p53*^-/-^ MEFs had no effect on LARP1 expression, Supplementary Figure 6. Given the dual role of LARP1 as both an enhancer and repressor of 5’-TOP mRNA translation^23^ it is possible that elevated expression of LARP1 in this context could drive 5’-TOP mRNA translation. However, knockdown of LARP1 had little to no effect on the expression of several 5’-TOP genes in *Arf^-/-^* MEFs, Figure 4g-h. Only one of the two shRNAs targeting LARP1 had a significant effect on expression but for only one of the 5’-TOP genes assessed – TPT1. These findings indicate that elevated LARP1 expression isn’t likely to be the driving force behind increased 5’-TOP mRNA translation in following loss of ARF, though it may have a role in the regulation of specific 5’-TOP mRNAs.

**Figure 5:**
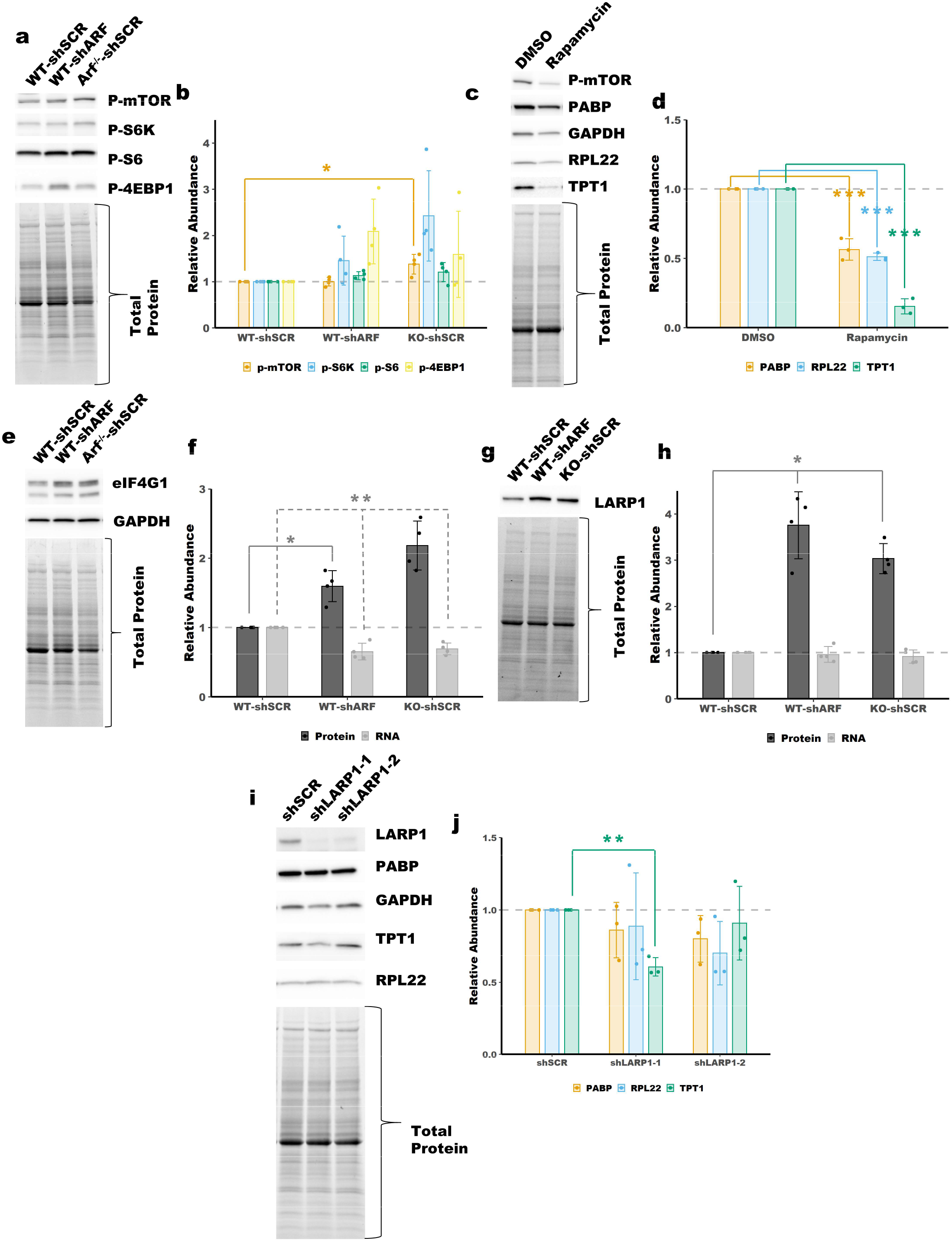
The expression and activity of 5’-TOP mRNA regulators following loss of tumor suppressors. **a** Immunoblot analysis showing activity of mTORC1 in WT and ARF-null MEFs and quantified in **b**. p-mTOR = p-S2484, p-S6K = p-T389 (p70), p-S6 = p-S240/S244, p-4EBP1 = p-T37/T46. Total protein was used for normalization. Mean±SD, n=4 **c** Immunoblot analysis showing expression of 5’-TOP mRNA encoded proteins following treatment with rapamycin (Selleck Chemicals) and quantified in **d**. *Af*^-/-^ MEFs were treated with 5 nM rapamycin for 4 days. Total protein was used for normalization. Mean±SD, n=3. **e** Immunoblot analysis showing increased expression of eIF4G1 in ARF-null MEFs. **f** Quantitation of blot in **e** and qPCR for eIF4G1, total protein was used for normalization of immunoblot, Mean±SD, n=4. **g** Immunoblot analysis showing increased expression of LARP1 in ARF-null MEFs. **h** Quantitation of blot in **g** and qPCR for LARP1, total protein was used for normalization of immunoblot, Mean±SD, n=4. **i** Immunoblot of *Arf*^-/-^ MEFs following knockdown of LARP1 with two different shRNAs. **j** Quantitiation of blot in **i**, total protein was used for normalization of immunoblot, Mean±SD, n=3. *, **, *** *p*-value < 0.05, <0.01, <0.001; t-test with Bonferoni correction.

### Loss of p53 causes upregulation of 5’-TOP mRNAs

Having observed a role for ARF in regulation 5’-TOP mRNA translation, we next sought to determine if p53, which is tightly coupled to ARF expression, has a similar role. Polysome profiling revealed increased translation in *p53*^-/-^ MEFs, as indicated by higher polysome peaks, Figure 6a. We used immunoblot analysis and qPCR to determine the protein and mRNA expression of some of the 5’-TOP genes studied above. We observed an increase in protein expression for two of the 5’-TOP genes (PABP and TPT1) in the *p53*^-/-^, there was no increase in RNA expression, Figure 6b-d. This is consistent with our findings from ARF-null MEFs. Activity of the RPL23a reporter was significantly increased in *p53*^-/-^ MEFs, though the magnitude was small. However, this experiment is confounded by a decrease in the expression of the SV40 driven reporter. This could arise from a difference in the stability of *Renilla* or firefly luciferase in p53-null vs WT MEFs. Immunoblot analysis of WT and *p53*^-/-^ MEFs revealed an increase in LARP1 expression, like that observed in ARF-null MEFs, Figure 6b and 6e. There was no increase in LARP1 mRNA expression in p53-null MEFs relative to WT MEFs, Figure 6e.

**Figure 6:**
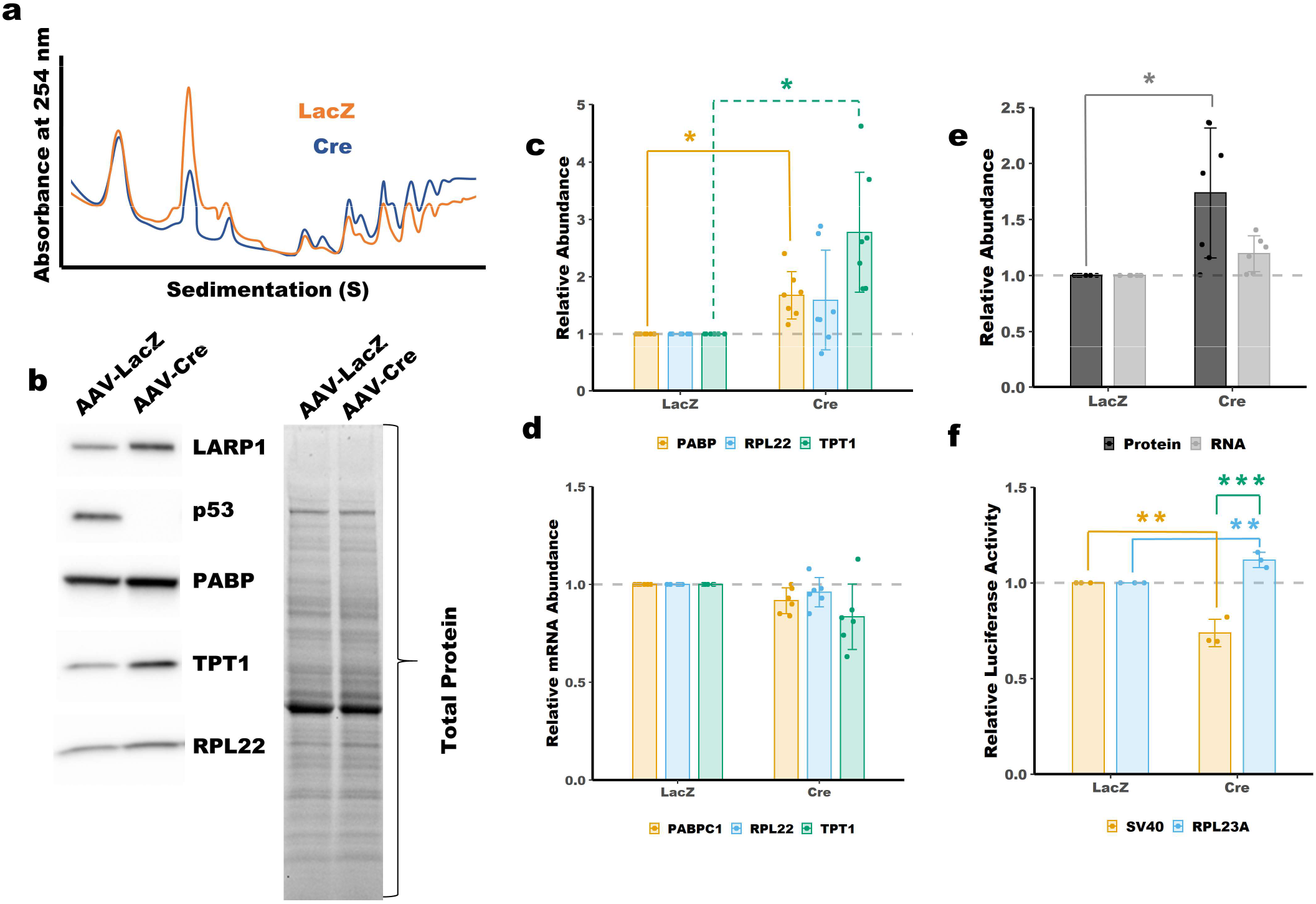
Upregulation of 5’-TOP mRNAs in *p53*^-/-^ MEFs. **a** Polysome profiling showing increased polysome peaks following knockout of *p53* (Cre). Immunoblot analysis showing increased protein abundance of LARP1, PABP, TPT1 and RPL22 following knockout of *p53* (AAV-Cre). **c** Quantitation of blots in panel **b**. Total protein was used for normalization. Mean±SD, n=7. **d** qPCR shows no change in mRNA expression of 5’-TOP mRNAs following knockout of *p53*. Mean±SD, n=6. **e** Quantitation of blot in **b** and qPCR for LARP1. Mean±SD, n=7(protein), 6(RNA). **f** Luciferase activity of a 5’-TOP reporter is increased in *p53*^-/-^ MEFs. Luciferase activity was normalized to *Renilla* luciferase transfection control and set relative to LacZ control. Mean±SD, n=3. *, **, *** *p*-value < 0.05, <0.01, <0.001; t-test with Bonferoni correction.

## Discussion

Together these data fit well with what is known about the functions of ARF in ribosome biogenesis. ARF is known to repress ribosome biogenesis by affecting rRNA transcription, processing and nuclear export ^7–16^. Changes in 5’-TOP mRNA translation upon ARF-loss would help to balance concomitant changes in rRNA production by providing increases in the necessary ribosome subunit proteins. Increased translation of 5’-TOP mRNAs following loss of the ARF tumor suppressor provides a missing mechanism for the increased rates of translation and ribosome biogenesis seen in many human cancers ^30^.

While many of the functions of ARF in regulating ribosome biogenesis are p53 independent, we show here that the regulation of 5’-TOP mRNAs by ARF requires p53 expression. Furthermore, we show that loss of p53 partially phenocopies the effects of ARF-loss on 5’-TOP mRNA translation. These data suggest that canonical ARF-MDM2-p53 axis plays an important role in 5’-TOP mRNA translation. Interestingly, free ribosomal proteins, such as RPL11, are known to activate p53 through binding MDM2^18^. Elevated expression 5’-TOP containing ribosomal proteins following p53-loss would complete a feedback loop.

It is unclear what factors drive increased 5’-TOP mRNA translation following loss of ARF. We observed increased expression and activity for several regulators of 5’-TOP mRNA translation. Increased mTORC1 activity likely contributes to improved translation efficiency of 5’-TOP mRNAs in ARF-null cells, however the increased mTORC1 activity observed in ARF-null MEFs is very modest. By inhibiting mTOR with rapamycin we showed that maximal expression of 5’-TOP encoded proteins requires mTOR activity. mTORC1 regulates 5’-TOP mRNA translation through phosphorylation and inactivation of LARP1. LARP1 acts to balance 5’-TOP mRNA translation through inhibition of translation when dephosphorylated and enhancing translation when phosphorylated ^23^. Dephosphorylated LARP1 binds the 5’-TOP motif and blocks binding of eIF4G1. We observed increased expression of both eIF4G1 and LARP1 following ARF-loss. Because of technical limitations described above it is difficult to assess the role of eIF4G1 via overexpression in this system. In addition, because eIF4G1 is a key component of translation initiation, knockdown of eIF4G1 would likely be cytotoxic or generate many pleiotropic effects making it difficult to evaluate the role of eIF4G1 expression in regulation 5’-TOP mRNA translation following loss of ARF. While we were unable to assess the role of eIF4G1 in this system we did investigate the importance of LARP1 expression. Knockdown of LARP1 in *Arf*^-/-^ MEFs had only modest effects on 5’-TOP mRNA expression, with one shRNA causing a significant reduction in the expression of the 5’-TOP mRNA encoded protein TPT1. This suggests that at least for TPT1, LARP1 may modestly enhance translation. Because phosphorylated LARP1 is known to enhance 5’-TOP mRNA translation this result is consistent with active mTORC1 in *Arf*^-/-^ MEFs. We suspect that increased 5’-TOP mRNA translation in ARF-null MEFs is likely caused not by increased activity or expression of mTORC1, eIF4G1 or LARP1 alone – but instead small changes in activity or expression of each of those factors contributed to increased 5’-TOP mRNA translation.

## Materials and Methods

### Cell culture

WT (C57BL/6J), Arf^-/-^ (B6.129X1-*Cdkn2^atm1Cjs^*/KaiJ), *p53^flox/flox^*(FVB.129-*Trp53^tm1Brn^*) mouse embryonic fibroblasts were cultured in Dulbecco’s modified Eagle’s medium (DMEM) (Hyclone) with 10% fetal bovine serum (Invitrogen), 2 mM glutamine (Hyclone), 0.1 mM nonessential amino acids (Hyclone), 1 mM sodium pyruvate (Hyclone), and 2 μg/ml gentamicin (Invitrogen). Low passage MEFs, between 1-4, were used for all experiments.

### Viral Production and Transduction

Lentivirus was produced by Lipofectamine 2000 (Inivitrogen) transfection of 293T cells with pCMV-VSV-G, pCMV-ΔR8.2, and either pLKO.1-puro for shRNAs or pLVX-puro for overexpression. Virus was harvested 48 hours post-transfection. Cells were transduced with lentivirus for 16 hours in the presence of 10 μg/mL protamine sulfate. The cells were selected with puromycin at 2 μg/mL for two days. The sequences for the shRNA-scramble (shSCR) and shRNA-ARF (shARF) were described previously ^16^.

### Measurement of Bulk Translation

For polysome profiling, translation was inhibited by the addition of 50 μg/mL cycloheximide for 5 minutes at 37 °C in culture media. The cells were then washed with PBS, trypsinized and resuspened in culture media. The cells were pelleted and washed with PBS prior to lysis in polysome lysis buffer (20 mM Tris pH 7.26, 130 mM KCl, 10 mM MgCl_2_, 2.5 mM DTT, 0.5% NP-40, 0.2 mg/mL heparin, 0.5% sodium deoxycholate, 50 μg/mL cycloheximide and 200 units/mL RNasin (Invitrogen)). Lysis occurred over 10 minutes on ice prior to clarification at 8,000 g for 10 minutes at 4 °C. The absorbance at 260 nm was determined for each sample and an equal number of absorbance units for each sample was overlaid onto a 10-50% sucrose gradient made with sucrose gradient buffer (10 mM Tris pH 7.26, 60 mM KCl, 10 mM MgCl_2_, 1 mM DTT, 0.1 mg/mL heparin, 10 μg/mL cycloheximide). The gradients were subjected to ultracentrifugation at 36000 rpm for 3 hours at 4 °C. A Teledyne Isco fractionation system with UV detector was used to determine absorbance at 254 nm along the gradient.

For measurement of translation rates by puromycin incorporation, MEFs were treated with 10 μM puromycin at room temperature for the times indicated. As a control, cells were also treated with 100 μg/mL cycloheximide for 5 minutes at 37 °C prior to addition of puromycin. Immediately following treatment, the cells were washed with 1x PBS containing 100 μg/mL cycloheximide to inhibit further puromycin labeling. The cells were harvested and lysed in RIPA buffer with 1x HALT (Pierce) and 100 μg/mL cycloheximide. Immunoblot was performed as described above with anti-puromycin antibody (DSHB, University of Iowa).

### Plasmid Construction

The 5’-TOP reporter for Rpl23a was made by PCR amplification of a region encompassing 500 bp upstream of the annotated transcription start site to the start codon using primers RPL23A Promoter Forward: 5’-GTACCTCGAGGAGCTATAAAGGGAAACCCTGTCTC −3’ and RPL23A 5’UTR Reverse: 5’-GTACCCATGGTGCI IGGCTGAAAAGGATGGCCC-3’. The PCR product was digested with XhoI and NcoI and ligated into pGL3-control (Promega). The resulting plasmid, pGL3-RPL23A-FF, was confirmed by Sanger sequencing.

To make pGL3-Renilla, Renilla luciferase was PCR amplified from pMT-DEST48-FLP ^31^ with Renilla Luciferase Forward: 5’-TGGAAGCTTGGCATTCCGGTACTGTTGGTAAAGCCACCATGACTTCGAAAGTTTATG-3’ and Renilla Luciferase Reverse: 5’-TGGAAGCTTTTATTATTGTTCATTTTTGAGAAC-3’ and digested with HindIII (NEB). Renilla luciferase was then ligated into pGL3-Control digested HindIII (NEB) to make pGL3-Renilla-FF. To remove the firefly luciferase coding sequence, the plasmid was digested with NarI and XbaI (NEB), subsequently the ends were blunted with Klenow (NEB) and the plasmid was ligated. The resulting plasmid pGL3-Renilla-deltaFF was confirmed by Sanger sequencing.

### Transfection of MEFs and Luciferase Assay

The day before transfection 1×10^5^ cells were plated per well in a six well dish. The cells were transfected with Fugene6 (Promega) and 1 ug each of either pGL3-control or pGL3-Rpl23a and pGL3-Renilla. After 24 hours the cells were washed briefly with 1x PBS prior to measurement of luciferase activity by Dual Luciferase Reporter Assay (Promega).

### Ribosome Profiling and RNAseq

For RNAseq, total RNA was isolated from the appropriate cells using the Direct-zol RNA MiniPrep (Zymo) with Trizol (Invitrogen) for initial cell lysis. Contaminating genomic DNA was removed by treatment with Turbo DNA-free Kit per the manufacturers protocol (Invitrogen). The RNA concentration was determined by Qubit RNA BR Assay (Invitrogen) and ribosomal RNA was removed using the RiboZero Gold rRNA Removal Kit (Illumina) per the manufacturers protocol.

The ribo-depleted RNA (10 μL) was then fragmented by adding one volume of 2x Fragmentation Buffer (0.5 M EDTA, 0.1 M Na_2_CO_3_, 0.1 M NaHCO_3_) and incubating at 95 °C for 20 minutes. Fragmentation was inhibited by the addition of 280 μL of Stop Solution (3 M NaOAc pH 5.5, 15 mg/mL GlycoBlue (Invitrogen)). The RNA was the precipitated by the addition of ethanol. The RNA was resuspended in 10 mM Tris pH 8.0 and resolved on a 15% acrylamide Urea-TBE gel (Bio-Rad). A region containing fragments between 17-34 nt (the same size as those isolated for ribosome profiling ^32^) was excised and extracted overnight in RNA Extraction buffer (0.3 M NaOAc pH 5.5, 1 mM EDTA and 0.25% Sodium dodecylsulfate). The extracted RNA was precipitated by ethanol precipitation and subsequently used for sequencing library preparation as described below.

Ribosome footprinting was performed as described previously ^32^,with the use of RNase T1 (Thermo Scientific) instead of RNase I for the footprinting step, see Supplementary Figure 2. The lysates were treated with 700 units of RNase T1 for 1 hour at room temperature with gentle mixing.

Following purification of the RNA footprints, sequencing library production was carried out for both the fragmented total RNA and ribosome footprints using the previously described protocol ^32^. The RNAseq and ribosome profiling libraries were multiplexed and sequenced on a HiSeq 3000 (Illumina) by the Washington University Genome Technology Access Center.

### Data and Code Availability

Raw sequencing reads are available at https://www.ncbi.nlm.nih.gov/geo (XXXXXXXX). Scripts used for analysis of differential expression and TE are available at GitHub (XXXXXXXX).

### Analysis of Sequencing Data

The ribosome profiling and RNAseq reads were processed to remove short reads, reads lacking the sequencing adapter and adapter only reads using *fastx_clipper*^33^. Indexes, adapter and other sequences introduced during library production were removed by *umi-tools* ^34^ and *cutadapt*^35^. Sequencing reads mapping to rRNA or ncRNAs were removed using *Bowtie2* ^36^. The processed reads were then aligned to the mouse genome (UCSC, mm10) using *TopHat* ^37^. Read counts of those aligning to the coding sequence of each gene was then determined by *HTSeq* ^38^. Differential translation efficiency was determined using *DESeq2* ^39^.

### Immunoblot

Cell pellets were lysed and sonicated in RIPA Buffer (50 mM Tris pH 7.4, 150 mM NaCl, 1% Triton X-100, 0.1% sodium dodecyl sulfate and 0.5% sodium deoxycholate) with 1x HALT Protease Inhibitor (Pierce). Seventy-five micrograms of protein lysate were resolved on 4-12% TGX Acrylamide Stain-Free gels (Bio-Rad). Stain-Free gels were imaged prior to transfer to PVDF membrane (Millipore). The blots were then probed with the appropriate primary antibodies: Primary antibodies: Abcam - PABP (ab21060); Bethyl - GAPDH (A300- 641A), LARP1 (A302-087A); Cell Signaling Technologies - TPT1(5128), p-MTOR(2971), p-S6(2215S), p-S6K(9205S), p-4EBP1(2855S), eIF4G1(2858), p53(2524S); Santa Cruz Biotechnology - ARF(sc-32748), RPL22(sc-136413), University of Iowa, DSHB - puromycin. Primary antibodies were detected with horseradish-peroxidase conjugated secondary antibodies (Jackson ImmunoResearch) and detection was carried out with Clarity Western ECL Substrate (Bio-Rad).

### Quantitative PCR

Total RNA was isolated using the Nucleospin RNA (Macherey-Nagel) with on column DNase treatment. Reverse transcription to make cDNA was performed with iScript Supermix (Bio-Rad). For qPCR the primers listed in the Supplementary Information were used with iQ Sybr Green (Bio-Rad). Fold change in RNA expression was determined by the ΔΔCt method with normalization to PSMA5, TOMM20 and ATP5B. The normalization genes were chosen based on steady expression across all samples as measured by RNA-seq.

## Acknowledgements

We thank Dr. Len Maggi for preparation of MEFs and guidance. This work was supported by: 5T32HL007088-42 and F32GM131514 to KAC; R01CA190986-03 and DOD W81XWH-15-1-0388 to JDW and the Longer Life Foundation: An RGA/Washington University Partnership.

## Author Contributions

KAC and JDW conceived the project. KAC performed the experiments and data analysis. RC generated the luciferase reporter constructs. KAC wrote the manuscript. All authors edited the manuscript.

