## Supplemental Information for "ARF suppresses 5’-terminal oligopyrimidine mRNA translation"

**Supplementary Information - ARF suppresses 5'-terminal oligopyrimidine mRNA translation**

**Quantitative PCR Primers (5' -> 3')**

PSMA5 qF
ATTGGCTCTGCTTCTGAGGG
PSMA5 qR
ATGAGCGAGGACTTGATGGC
TOMM20 qF
TTACAGCAGACTCTTCCGCC
TOMM20 qR
TCTTCAGCCAAGCTCTGAGC
ATP5B qF
TGCAGGAAAGGATCACCACC
ATP5B qR
AATAGCCCGGGACAACACAG
Renilla-qF
ATAACTGGTCCGCAGTGGTG
Renilla-qR
TAAGAAGAGGCCGCGTTACC
Firefly-qF
TCCATCTTGCTCCAACACCC
Firefly-qR
TCATCGTCTTTCCGTGCTCC
LARP1 qF
TCCTTTCACCGTGTACAGGC
LARP1 qR
TGCGGACTTTCTCCTCAACC
RPL22 qF
TGAAGTTCACCCTGGACTGC
RPL22 qR
GGTTGCCAGCTTTCCCATTC
PABPC1 qF
GTTACCATGCAACAGCCTGC
PABPC1 qR
GAAACAGCCGTTCACCCAAC
TPT1 qF
AGCCATGACGAGCTGTTCTC
TPT1 qR
TTTCCACCGATGAGCGAGTC

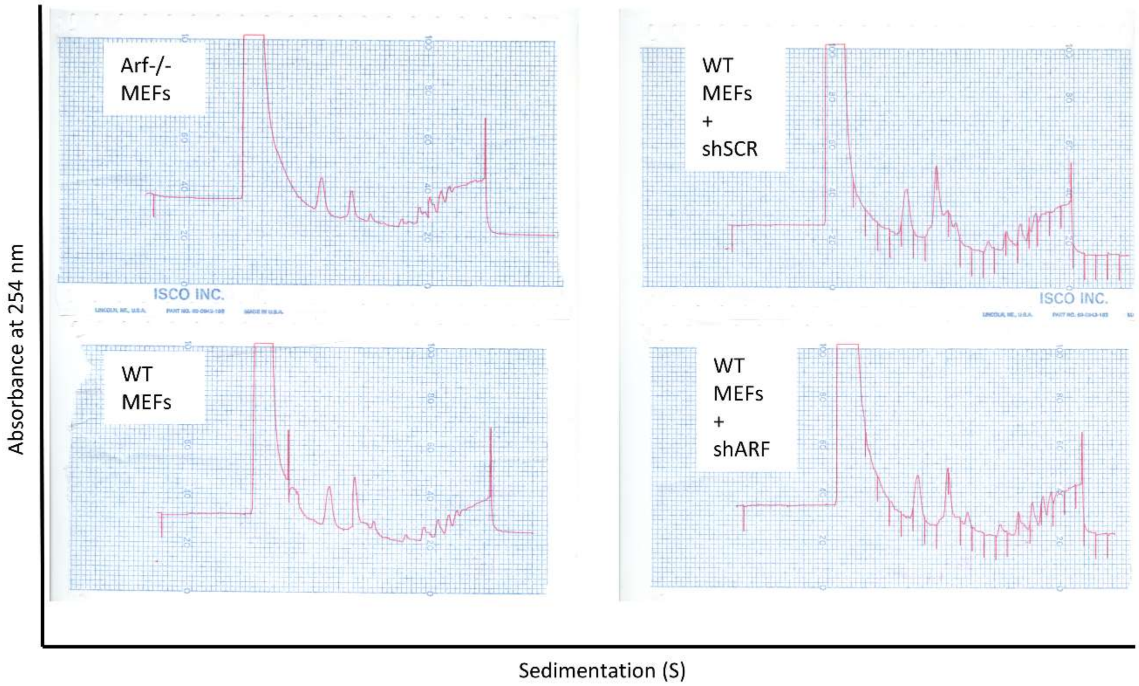


**Supplementary Figure 1: Raw polysome profiling traces for ARF knockout and knockdown MEFs**

The analog polysome profiling traces on the left were digitally traced in Microsoft PowerPoint to make panel **a** in **Figure 1** of the main text. The sudden spike in the WT MEFs trace corresponds to a bubble passing through the detector.

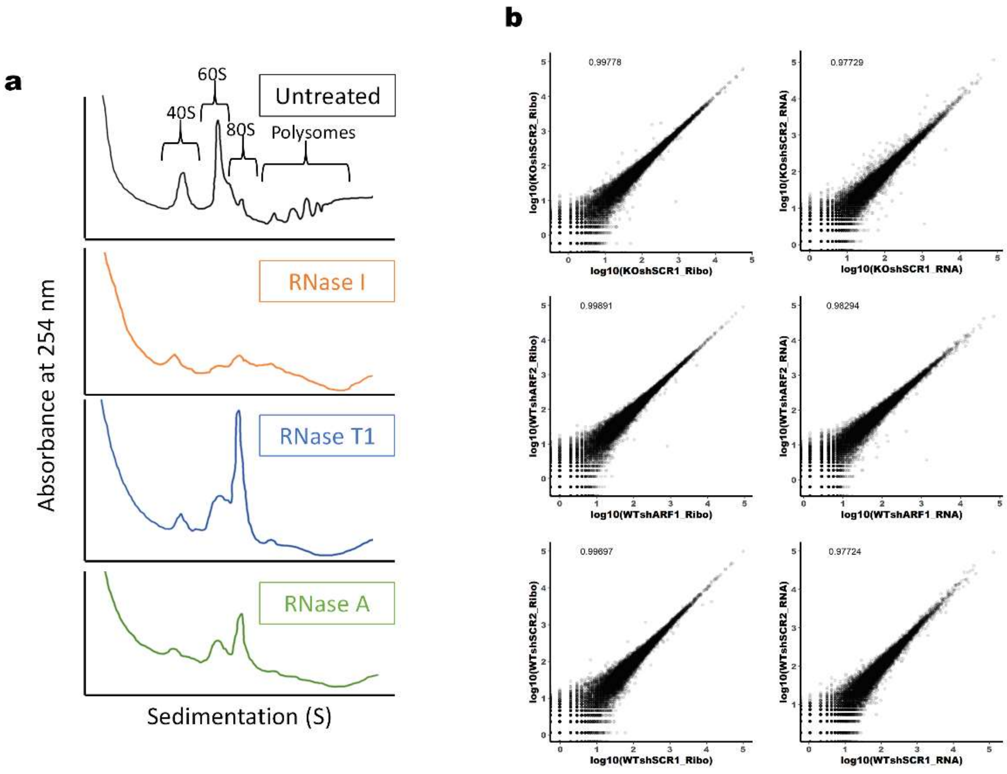


**Supplementary Figure 2: Ribosome profiling nuclease selection and reproducibility**

Panel **a** shows the effect of ribonuclease treatment of MEF lysates using RNase I, T1 or A. RNase T1 completely removed polysome peaks while maintaining a strong monosome peak. **b** Replicate-to-replicate comparisons of ribosome profiling and RNAseq normalized read counts per gene.

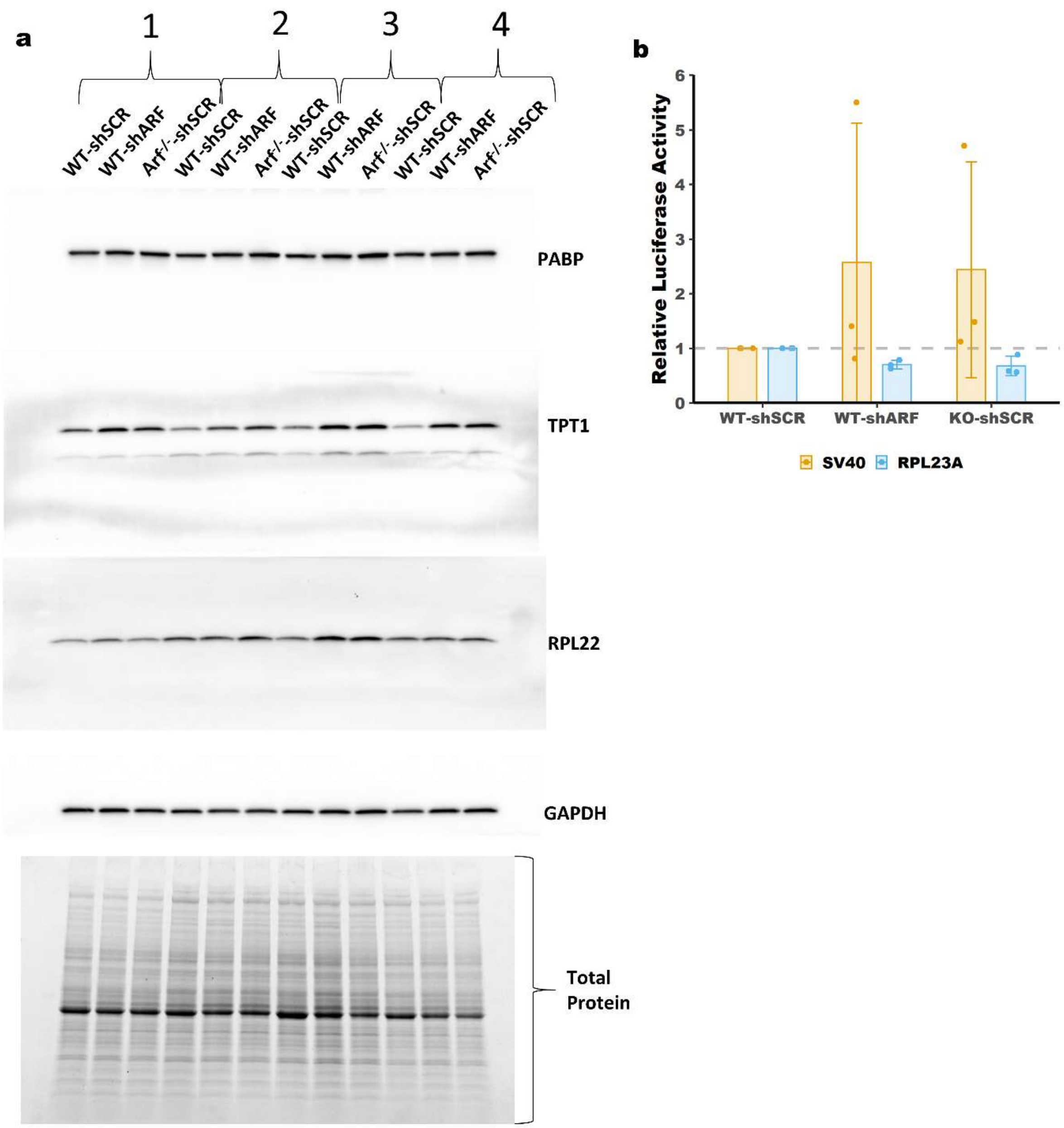


**Supplementary Figure 3: Extended data for Figure 3**

**a** Uncropped immunoblots for **Figure 3**. **b** qPCR of luciferase reporters following knockdown or knockout of ARF.

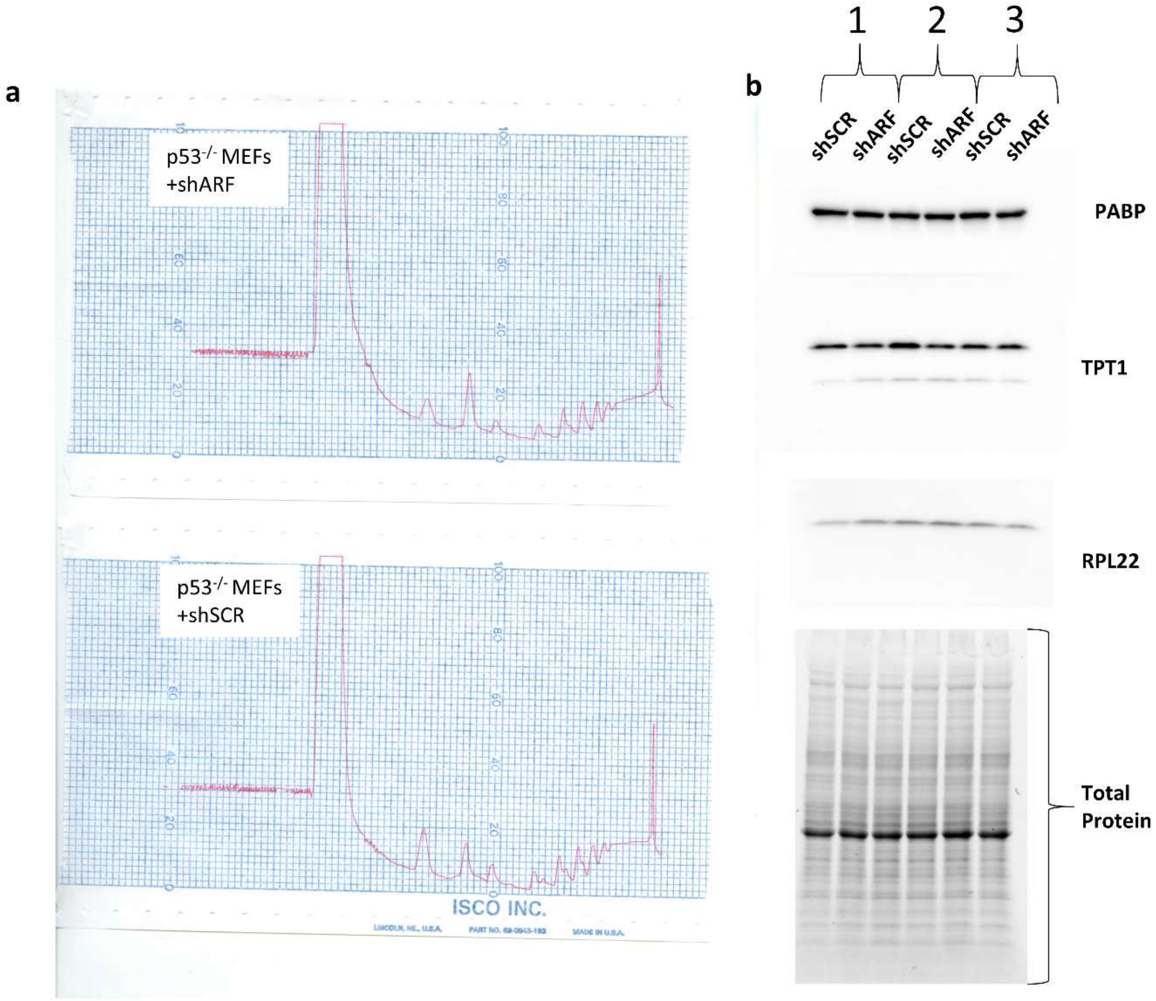


**Supplementary Figure 4: Source images for Figure 4**

Analog polysome profiling trace used in **Figure 4** and uncropped immunoblots. See main text for further details.

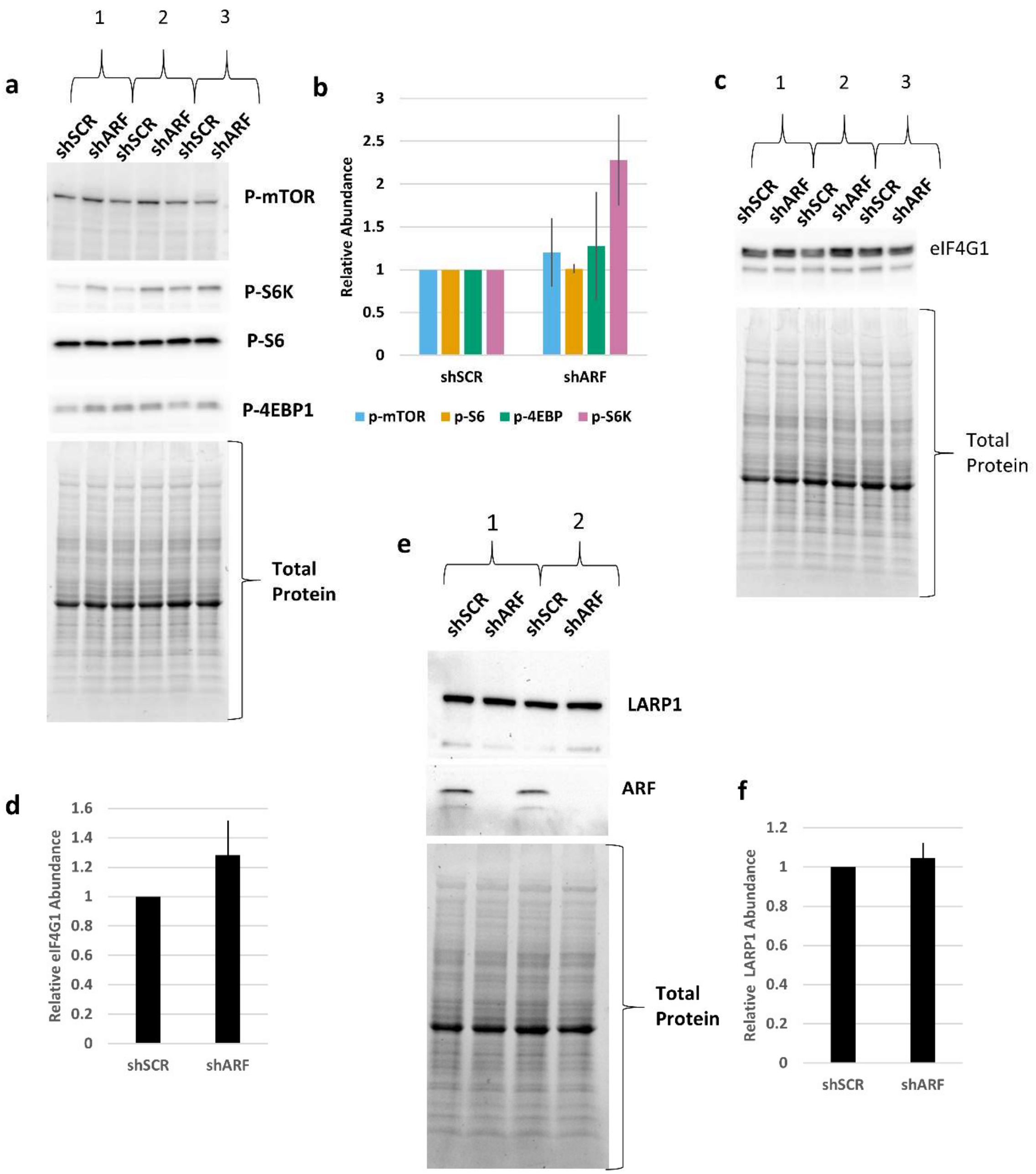


**Supplementary Figure 5: Analysis of 5'-TOP regulators expression and activity following knockdown of ARF in *p53*$^{-/-}$ MEFs**

**a** assessment of mTORC1 activity following knockdown of ARF in *p53*$^{-/-}$ MEFs. **b** Quantitation of the immunoblot shown in panel **a**. Mean +/- standard deviation, N = 3. **c** assessment of eIF4G1 expression following knockdown of ARF in *p53*$^{-/-}$ MEFs. **d** Quantitation of the immunoblot shown in panel **c**. Mean +/- standard deviation, N = 3. **e** assessment of LARP1 expression following knockdown of ARF in *p53*$^{-/-}$ MEFs. **f** Quantitation of the immunoblot shown in panel **e**, Mean +/- standard deviation, N = 2.

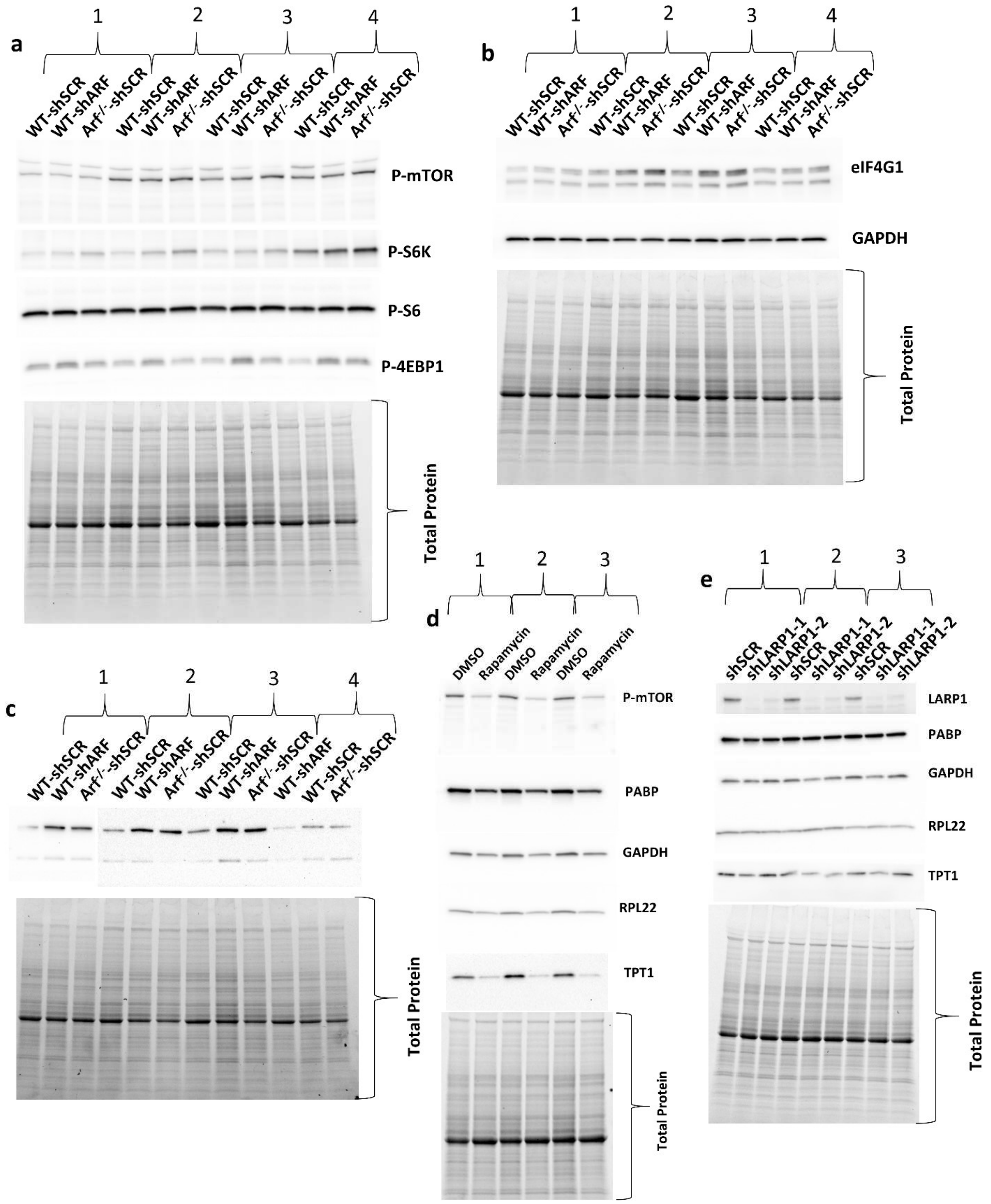


**Supplementary Figure 6: Source Images for Figure 5**

Uncropped immunoblots from **Figure 5**. See main text for further details.

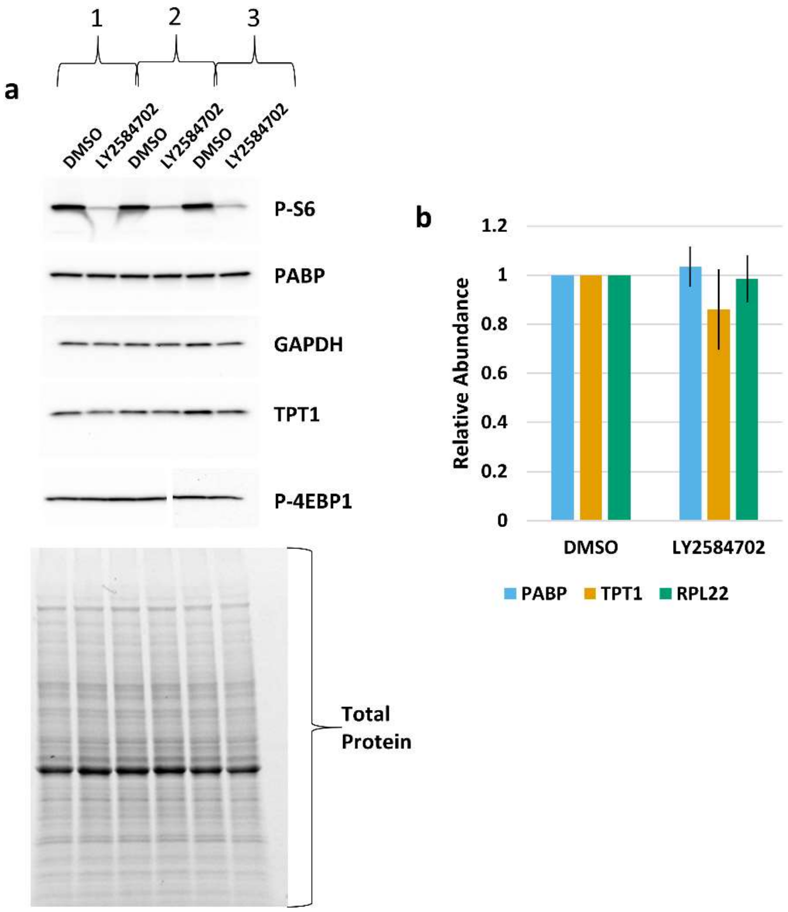


**Supplementary Figure 7: S6-Kinase inhibition has no effect on 5'-TOP mRNA expression**

*Arf$^{-/-}$* MEFs were treated with LY2584702 (ApexBio Technology) at a concentration of 1 µM for 4 days prior to harvesting. **a** Immunoblot analysis of 5'-TOP mRNA encoded proteins and P-S6 following treatment, quantified in panel **b**. Mean +/- standard deviation, n = 3.

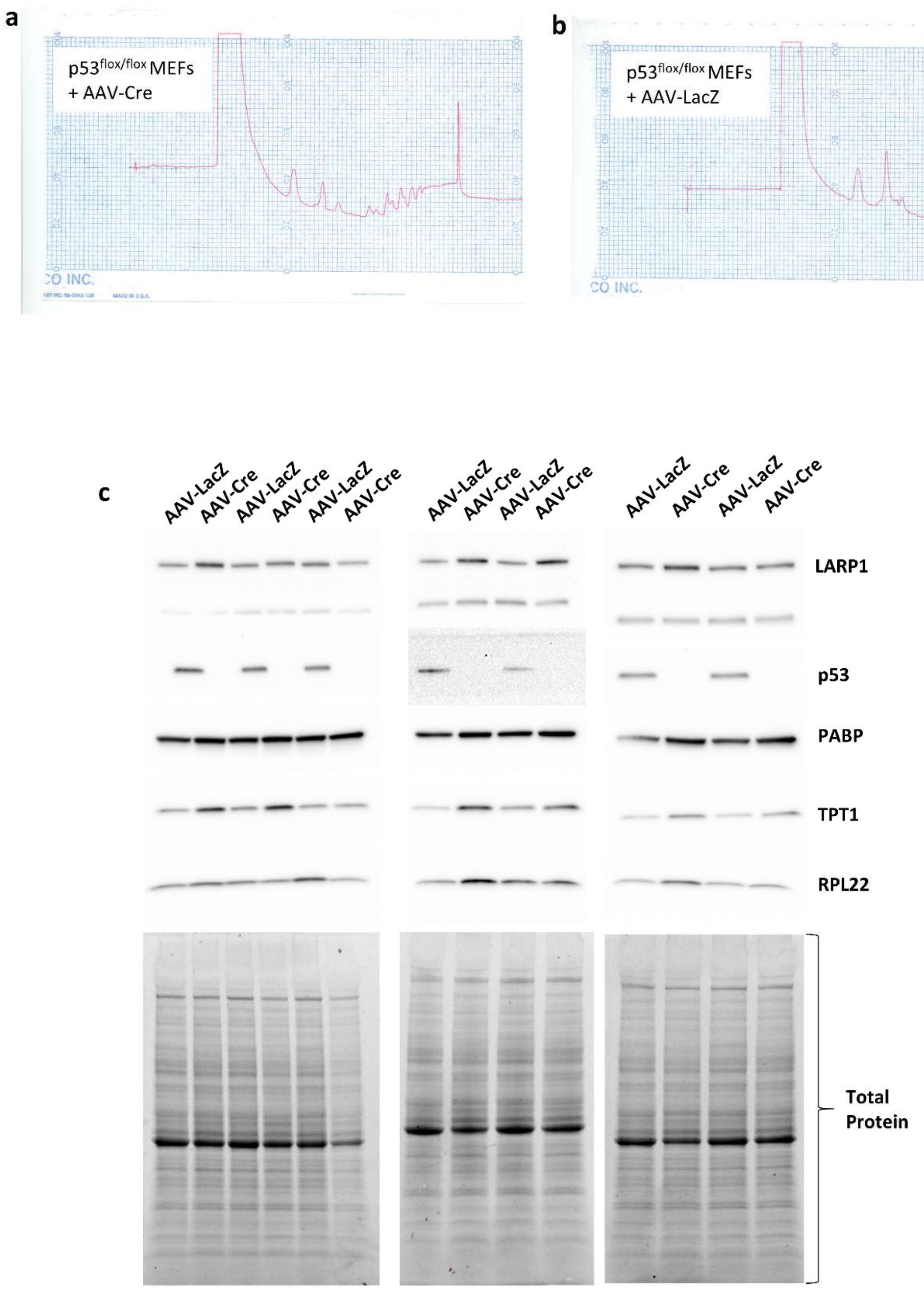


**Supplementary Figure 8: Source images for Figure 6**

Analog trace used in **Figure 6** and uncropped immunoblots. See main text for further details.
